# A new genetically engineered transplant model of glioma recapitulates key phenotypes of low- and high-grade gliomas

**DOI:** 10.1101/2025.10.29.685449

**Authors:** Devlin R. Forsythe, Shannon J. Oliver, Adam Valkovic, Danielle Lewthwaite, Germaine Uys, Brittany Koning, David Eccles, Laveniya Satgunaseelan, Ian. F. Hermans, Saskia Freytag, Jim R Whittle, Sarah A. Best, Melanie J. McConnell

**Author notes:** **Corresponding author:** Associate Professor Melanie-Jane McConnell, School of Biological Sciences, Te Herenga Waka Victoria University of Wellington. 7 Kelburn Parade, Kelburn Wellington 6012, New Zealand.

## Abstract

Immune competent animal models are essential in preclinical glioma research. The ability to investigate tumor development with key tumor microenvironment components such as immune infiltration, stromal cells and extracellular matrix allows for the investigation of these complex tumors. However, the current range of syngeneic models of glioma possess intrinsic limitations which must be acknowledged when designing a preclinical study. To address this gap, we developed genetically engineered mouse cell line models (GEM-CLeMs) by introducing common glioma driver mutations into immortalized astrocytes. A high-grade glioma model was generated by combining *Pten* knockdown with *RAS* V12 overexpression, while a low-grade glioma model was produced through *p53* knockdown with mutant *IDH1*^R132H^ overexpression. The RAS/Pten GEM-CLeM tumors grew rapidly *in vivo*, displayed necrosis, multinucleated pleomorphisms, abundant vascularization, and showed strong enrichment for extracellular matrix remodelling and mesenchymal genes, features closely aligned with human glioblastoma. Importantly, the RAS/Pten GEM-CLeM tumors showed similar survival trends and immune infiltration patterns as a corresponding *Kras*^G12D^/*Pten*^cKO^ GEMM, but could be grown to larger sizes, facilitating better stromal and immune analyses. In contrast, the *IDH1*^R132H^/p53 GEM-CLeM formed slow-growing tumors with distinctive immune infiltration and vascular patterns, consistent with low-grade glioma phenotypes. Compared with the commonly used cell line GL261, the GEM-CLeM tumors had higher levels of stromal integration and immune suppression, making them a more faithful model of the glioma tumor microenvironment. This system enables rapid generation of transplantable glioma models with defined driver mutations in a low-mutational background, offering a flexible platform for dissecting glioma biology and evaluating immunotherapies.

**Importance of the study:** We have generated a set of customizable, modular Genetically Engineered Mouse Cell Line Models (GEM-CLeMs) of “high grade” and “low grade” glioma. They reliably form tumors when transplanted intracranially into immune-competent C57BL/6 mice, and they are cost- and time-effective at capturing the important characteristics of glioma, both mutant IDH1 low grade glioma and high grade glioblastoma. Critically these characteristics include the myeloid-rich immune suppressive tumor immune microenvironment, a key weakness of existing murine glioma cell lines like GL261.

These GEM-CLeM models can be used in multiple ways. The cells are amenable to further manipulation, so therapeutic targets and drug mechanism of action can be assessed. The activity of candidate genes in tumour formation and phenotype can be determined. Most importantly the models can be used to develop effective immunotherapies, including strategies to target macrophage and myeloid cell immune suppression.

**Key points:**

- Combinations of driver mutations were engineered into an immortalised mouse astrocyte.
- Engineered cells formed tumours on intracranial transplant into immune competent mice.
- Tumors had key histological and immune suppressive features of human glioma.

## Introduction

Gliomas range from the slow growing low-grade glioma (LGG) to the highly malignant glioblastoma (GBM). The presentations of glioma vary substantially histologically, likely due to the different glial cells of origin, such as neural stem cells (NSCs), oligodendrocyte precursor cells (OPCs) and astrocytes^1^. The mutational status of the tumors also impacts presentation and growth. LGG are defined by mutation in *IDH1/2*, with frequent alterations in *TP53* at the early stages of gliomagenesis^2^ and grade progression associated with *CDKN2A/B* alterations^3^.

The aggressive GBM are driven by the accumulation of multiple driver mutations which have been identified by extensive genome sequencing^4-6^. Mutations to the MAPK and AKT/mTOR pathways, such as *EGFR* amplification and mutation^4,6^ or *PTEN* deletion (often as part of chromosome 10 loss)^4,7^, are common drivers of GBM which contribute to rapid proliferation and high levels of invasion into surrounding healthy tissue^8^. However, selective targeting of oncogene drivers like EGFR has had little to no impact on the treatment of GBM. Furthermore, GBM commonly have an immunologically “cold” microenvironment induced by an influx of immune suppressive myeloid cells^9,10^, and have not yet benefitted from immunotherapy, because this myeloid suppression has not yet been addressed.

This culminates in extremely poor survival for patients with GBM. Despite a well-established standard of care - maximally safe resection of the tumor followed by adjuvant temozolomide and radiation therapy^11^-the median survival for patients is still 15 months, with minimal improvement over the past two decades^12^. While there are many reasons for this lack of progress, such as intrinsic resistance to apoptosis^13,14^and the blood-brain barrier^15,16^, the absence of appropriate preclinical animal models has also hindered the translation of many new treatments^17^.

Patient derived xenografts (PDX) are frequently used for modelling GBM in mice^18^. Transplanted human tissue maintains the genomic heterogeneity that characterizes GBM, however, PDX transplants require severely immunosuppressed mice for the xenograft to grow^19^. This lack of immune integration into the tumor removes a significant component and key characteristic of human GBM^20^. To counter this, mouse glioma cells such as GL261^21^ and CT2A^22^ can be used in immune competent mice. These murine cell lines rapidly and reliable form tumors, which makes them the most commonly used when fully immune competent mice are required^23^. However, these tumors miss key GBM characteristics, such as the infiltrative advancing edge^24^ and an immune suppressive microenvironment^25^. Furthermore, both syngeneic and PDX transplant models have failed to faithfully recapitulate the *IDH1* mutation both when cultured *in vitro* and transplanted *in vivo*^26^, which significantly limits the ability of researchers to accurately study LGG.

Genetically engineered mouse models (GEMMs) are an alternative which can address these limitations^27^. Tumors formed by mutations *in vivo*, either through viral transduction or conditional knockouts, provide a useful tool to investigate how specific mutations contribute to glioma development. The use of GEMMs has also enabled *IDH1* mutations to be modelled in mice - through the introduction of *Idh1*^R132H^ construct into cells in the sub ventricular zone, tumors were induced by expression of the mutant enzyme^28^. However, GEMM tumors can be variable in the time taken to form a tumor, and they can take months to form. This is slow compared with transplantation models which grow in days to weeks^17^. Additionally, development and maintenance of the mouse strains required to generate new GEMM is time and resource-intensive. Overall, the absence of an efficient, cost-effective way to generate new immune-competent preclinical model is a significant gap in the glioma research field.

To address these limitations, we developed a new approach to preclinical glioma models. Using an immortalized C57BL/6 astrocyte cell line^29^, mutations specific to human glioma were introduced to generate a new transplantable glioma model. The resulting genetically engineered cell line models (GEM-CLeM) are easily modified to express different combinations of genetic drivers, and grow in immune competent mice. Here, we created models of GBM and *IDH1*-mutant LGG and characterized how faithfully they recapitulated salient features of human glioma.

## Methods

The eGFP parental astrocyte cell line was generated from the cortex and hippocampus of neonatal C57BL/6 eGFP transgenic mice^29^. The murine cell line GL261 was purchased from Fredrick National Laboratory (MD USA). Cells were cultured in DMEM high glucose (ThermoFisher Scientific, NZ) supplemented with 10% FBS and grown at 37°C with 5% CO_2_. Cells were routinely tested for mycoplasma using PCR.

pX330 CRISPR plasmids targeting murine Pten and P53 were published^30^. Parental astrocytes were transfected with 500ng pX330 plasmid, 500ng tdTomato reporter plasmid and 3µL ViaFect reagent (Promega). After 48 hours, eGFP+tdTomato+ single cells were FACS sorted, individual colonies expanded, then screened for CRISPR-induced knockdown with a Surveyor® endonuclease assay for heteroduplex formation. CRISPR-induced deletion and off-target mutations were assessed with whole genome sequencing (Grafton Clinical Genomics, New Zealand). Paired-end reads were mapped to GRCm39/mm10 and visualized on IGV. Potential off-target sites were predicted with TagScan, scored^31^, and visualized on IGV.

Pten and *p53* knockdowns were confirmed by western blot, with p53KD cells irradiated with 2Gy to stabilize *p53* 2 hours prior to lysis. Cells were lysed (70 mM NaCl, 20 mM Tris-HCL pH 7.4, 0.1% IGEPAL (v/v)) and 30µg of lysate separated on 12% SDS-PAGE. Proteins were transferred to PVDF, blocked with 5% BSA and incubated with primary antibodies diluted 1:1000 with 5% BSA (Pten: Cell Signalling, 95595; P53: AbCam_PAb240). Secondary-HRP conjugated antibodies diluted 1:20,000 in 0.05% PBS/Triton X-100 (αRabbit: AbCam_150166; αMouse: BioLegend_405306) were incubated 1 hour, and detected using enhanced chemiluminescence (ThermoFisher Scientific, NZ).

Pten and *p53* knockdown lines were transfected with pcDNA3-H-Ras_V12^32^ and pcDNA3-Flag-IDH1-R132H^33^ and stable expression selected with 500 µg/mL G418 (ThermoFisher Scientific, NZ). Stable transfections were confirmed by immunofluorescence for RAS V12 (Cell Signaling Technology, NZ) and IDH1^R132H^ (Merck, NZ), and Goat-αIgG AF647 (ThermoFisher Scientific. αRabbit_A-21245. αMouse_A-21245). Nuclei were stained with DAPI (ThermoFisher Scientific, NZ) and fluorescence visualized by confocal microscopy (FV3000, Olympus NZ) with 405nm and 640nm lasers.

Cell proliferation was quantified (Incucyte S3, Sartorius, Germany). Cells plated at 5% confluence were monitored over 96 hours. Cell migration was assessed by wound healing – cells at 90% confluence in 96 well plate were uniformly scratched (WoundMaker96™) and media replaced with FBS-free DMEM to inhibit proliferation. Wound density was quantified over 12 hours (Incucyte S3).

C57BL/6 mice were purchased from The Jackson Laboratory (Bar Harbor, ME, USA) and bred and housed at the Biomedical Research Unit at Malaghan Institute of Medical Research. Male and female mice were maintained at 21°C with 45-65% humidity and 12-hour day-night cycles, and used for intra-cranial transplant at 10-12 weeks. Animal experiments were conducted under approvals from Victoria University of Wellington Animal Ethics Committee (AEC30425 and AEC30735), and the Walter and Eliza Hall Institute of Medical Research (AEC_2022.004).

Kras^G12D^ (B6.129S4-*Kras*^tmTyj^/J), Pten^floxed^ (B6.129S4-*Pten*^tm1Hwu^/J) and LSL-luciferase^T/+^ (Gt(ROSA)26Sortm1(Luc)Kael/J) alleles were obtained from The Jackson Laboratory. Mouse lines were intercrossed and maintained on a pure C57BL/6 background. Gene recombination was initiated by intra-cerebroventricular injection of postnatal day 2 pups with 2µL of 3×10^10^ PFU/mL Ad5-CMV-Cre (University of Iowa Gene Transfer Core Facility #VVC-U of Iowa-5). Tumor development was monitored by in vivo imaging (IVIS, Revvity) following a 100µL intraperitoneal injection of 10mg/mL D-luciferin (Revvity #122799).

Mice were anaesthetized with intraperitoneal ketamine/xylazine, then 2.5×10^4^ cells in 2µL PBS were transplanted intracranially (2 mm lateral, 1 mm posterior to bregma, depth 3 mm)^24^ with a 10µL 27 G Hamilton syringe and stereotactic head frame (Stoelting, IL, USA). Carprofen and buprenorphine were administered subcutaneously as pain relief pre-surgery, followed by carprofen post-surgery.

Mice were monitored for symptoms and euthanized either when pain was detected (grimace scale), 10% decrease in weight from transplant date, 15% decrease in weight over 48-hours, or onset of neurological symptoms. Mice were euthanized by CO_2_ asphyxiation, then cardiac perfused with PBS. Tumor-bearing hemispheres were fixed in 4% PFA for 18 hours then immersed in 70% ethanol, or cryopreserved fresh in OCT with liquid nitrogen. PFA-fixed tissue was paraffin embedded, sectioned to 5µm, and adhered to the slide at 50°C. Cryopreserved tissue was sectioned to 10µm and adhered to the slides at room temperature overnight.

For H&E staining and immunohistochemistry, tissue sections were rehydrated in 3x 100% xylene, 1x 100% ethanol, 1x 95% ethanol, then 1x 70% ethanol. Sections were stained with haematoxylin (Australian Biostain, 029-AHH) for 3 minutes, de-blued in 2% NH4 for 1 minute, and stained in eosin (Australian Biostain, 029-AEY) for 1 minute. Sections were dehydrated in 1x 70% ethanol, 1x 95% ethanol, 1x 100% ethanol, and 3x 100% xylene. Coverslides were mounted with DPX and adhered at RT overnight. H&E sections were visualized (VS200 Slide Scanner, Olympus, NZ and QuPath v0.4.4). Alternatively, slides were dewaxed in xylene, re-hydrated and antigen retrieval (1 mM EDTA pH 8, 0.0005% Tween20 in PBS) carried out in a steamer (32 minutes). Endogenous peroxidase activity was quenched using 3% Hydrogen Peroxide, 5 min. Sections were blocked in 5% goat serum and washed in 0.05% Tween20/PBS. CD45 primary antibody (Biolegend #103114 1/500 dilution) was incubated humidified for one hour. Biotinylated secondary antibodies and Vectastain Elite ABC HRP reagent (Vector Laboratories PK-7100) were each incubated for 30 min at RT. ImmPACT DAB Peroxidase substrate (Vector Laboratories SK-4105) was used according to manufacturer’s instructions, sections counterstained (15 sec Hematoxylin, 30 sec Scott’s tap water) and imaged (Nikon Eclipse 50i microscope, Axiovision software (Zeiss)).

For immunofluorescence, cryosections were fixed in 4% PFA for five minutes, then briefly washed in PBS to dissolve OCT. FFPE sections were rehydrated by immersion in 3x 100% xylene, 1x 100% ethanol, 1x 95% ethanol, and 1x 70% ethanol washes, then incubated for 30 minutes in high pH antigen retrieval buffer (eBioscience, 00-4956-58) at 80°C. All staining was conducted in a humidity chamber. Tissue was blocked (αCD16/CD32, 60 minutes) and immunolabelled for either 90 minutes at room temperature (cryosection) or overnight at 4°C (FFPE section), with the following primary antibodies diluted in PBS: CD45-PE (1:400, BioLegend_368510), CD31-AF594 (1:200, BioLegend_102520), CD105-BV421 (1:200, BD Biosciences_562760). Slides were washed in PBS before staining with either DAPI for five minutes (1:20,000, ThermoFisher Scientific, NZ) or DRAQ7 for 10 minutes (1:1000, Abcam, NZ). Coverslips were mounted (Agilent_S3023) and tissue visualized (FV3000 confocal microscope, Olympus, NZ)

RNA was extracted from endpoint tumor-bearing brains. Tumor and immediately adjacent tissue was dissected, homogenized, then snap frozen in liquid nitrogen. RNA was extracted (QIAzol tissue lysis reagent, Qiagen) chloroform extracted, and purified (Quick RNA MiniPrep, Zymo Research). RNA was quantified by fluorescence and spectrophotometry, and RNA integrity confirmed with TapeStation (Agilent, CA, USA).

Paired-end RNA sequencing (Novaseq 6000, Auckland Genomics, University of Auckland, NZ). Raw sequence data was processed with MultiQC (v1.25.1) Trimmomatic (v0.39), then aligned to M35 mouse transcript assembly with Salmon (v1.10.0). Differential expression analysis was performed with the DESeq2 R package (v3.20). Gene ontology analysis was conducted with clusterProfiler and gseGO^34^. Gene enrichment analysis was performed with fgsea utilising MSigDB gene sets^35^.

## Results

### Generation of high-grade glioma GEM-CLeM lines

An immortalized astrocyte-like cell line (Parental)^29^, was engineered with mutations frequently identified in glioma patients. These were chosen from published genomic data^4,6,36^ and have been validated as drivers in multiple GEMM studies^27,28,37^. To model an aggressive high-grade GBM, the tumor suppressor *Pten* was knocked down and *RAS* V12 oncogene over-expressed (Figure 1A).

**Figure 1.**
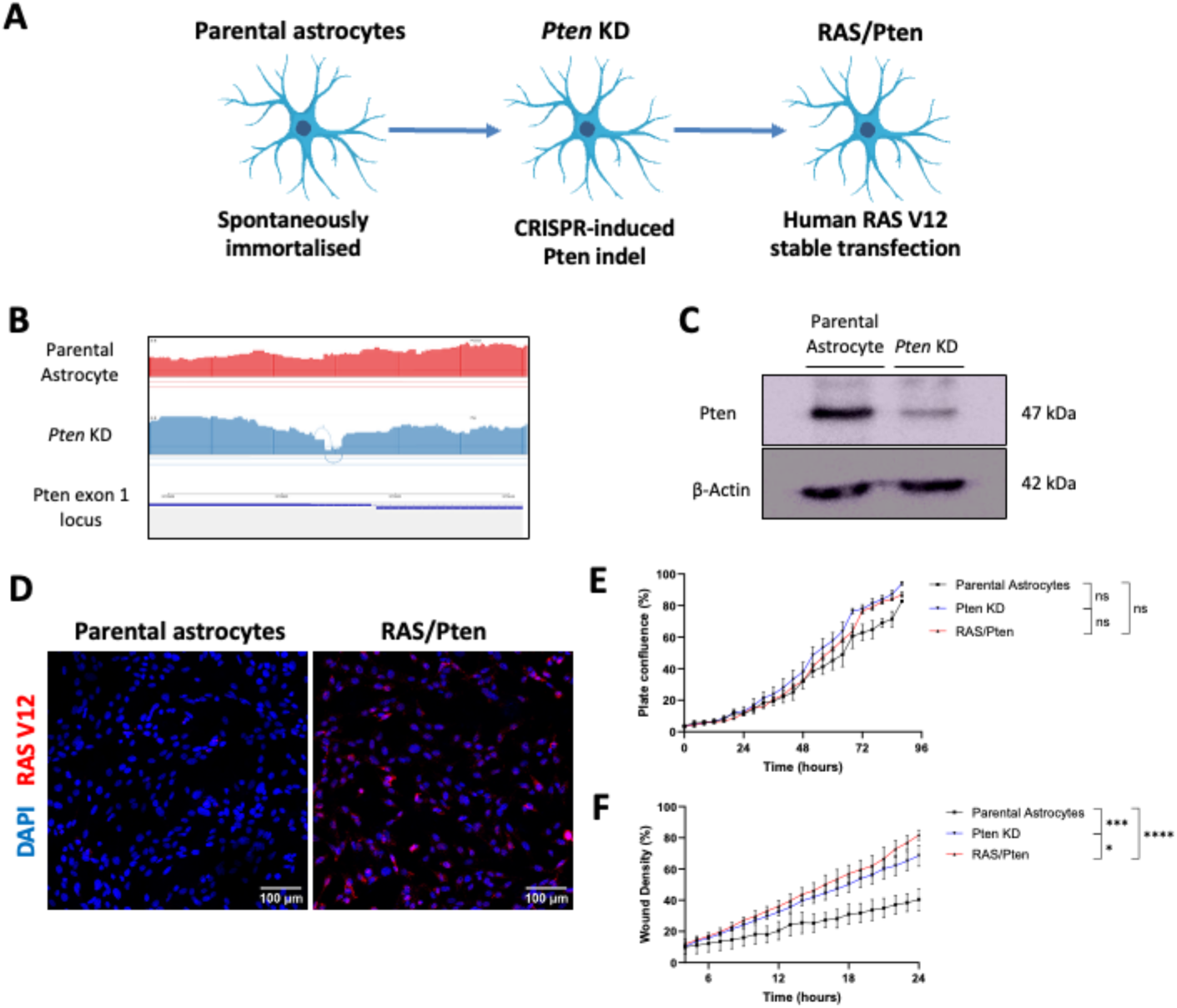
Development of the RAS/Pten GEM-CLeM cell line. A) Schematic showing the design with in vitro mutations to create *Pten* KD and *RAS/Pten* GEM-CLeM cell lines. B) Sashimi plot for WGS read depth over CRISPR sgRNA sites for Exon 1 of *Pten* in the Parental astrocytes (red) and Pten KD (blue) sequences. C) Western blot for Pten and B-actin protein expression, with parental astrocytes and Pten KD protein lysates. Blots were blotted for Pten, then stripped and reprobed for β-actin. D) Immunofluorescence for human RAS V12 protein expression (red) in the parental astrocytes and RAS/Pten GEM-CLeM cell line. Nuclei were stained with DAPI (blue). Scale bar, 100 microns. E) Proliferation rate of parental astrocytes (black), Pten KD (blue) and RAS/Pten (red) lines. F) Cellular migration rate of parental astrocytes (black), Pten KD (blue) and RAS/Pten (red) lines. Student t-test, * p<0.05, *** p = 0.005-0.001, **** < 0.001. ns = not significant. Error bars represent SEM and n = 3 biological replicates.

*Pten* indel mutations generated by CRISPR/Cas9 were screened by heteroduplex formation (Supplementary Figure S1) and confirmed by WGS (Figure 1B). Predicted off-target sites showed no change in sequence (Supplementary Figure S2). Consequent loss of Pten protein in the modified cells was demonstrated (Figure 1C) and loss of Pten function confirmed by increased Akt phosphorylation (Supplementary Figure S3). Next, overexpression of the human RAS V12 was confirmed using RT-PCR (Supplementary Figure S4) and the expression and subcellular localisation of RAS V12 was validated (Figure 1D). Thus, a genetically engineered mouse cell line model (GEM-CLeM) of *Pten* knockdown with RAS V12 gain (RAS/Pten) was generated.

The impact of the mutations on growth was determined *in vitro*. While not significant, *Pten* KD alone slightly increased the growth rate compared to the parental line (Figure 1E). No additional increase in growth rate was observed with introduction of RAS V12 (Figure 1E). Next, migration was investigated using a scratch wound healing assay. Compared with the parental cells, the *Pten* KD line showed a significant increase in migration, which increased further with the addition of RAS V12 (Figure 1F). These *in vitro* data suggest that the introduced oncogenic mutations induced an invasive phenotype in the parental astrocytes, consistent with human glioblastoma and other human GBM cell lines^8^.

### Tumorigenicity of RAS/Pten GEM-CLeMs

To investigate tumor-forming ability *in vivo*, the Pten KD and RAS/Pten lines were transplanted intracranially into immunocompetent C57BL/6 mice. A small number of cells (2.5×10^4^) were implanted and tumor lesions were allowed to develop until either tumor burden required the mice to be culled, or for 90 days, whichever came first. This was compared to the murine cell line GL261, a widely used intracranial transplant model (Figure 2A). As another comparator, the conditional alleles were introduced into neonate mice to create *Kras*^G12D^/*Pten*^cKO^/Luciferase (Luc)-reporter GEMM. Intracerebroventricular injection of a Cre-recombinase containing adenovirus (Ad5-CMV-Cre) was carried out at postnatal day 2 and monitored over time. Three different genotypes were produced through intracerebroventricular injection: *Kras*^G12D^/*Pten*^cKO^/Luc (n=4), *Kras*^G12D^/*Pten*^f/+^/Luc (n=1), and *Pten*^cKO^/Luc (n=2). While low numbers were used for this study, the GEMMs generated by *in vivo* mutagenesis had the same mutations and are directly comparable to the GEM-CLeM (Figure 2B).

**Figure 2.**
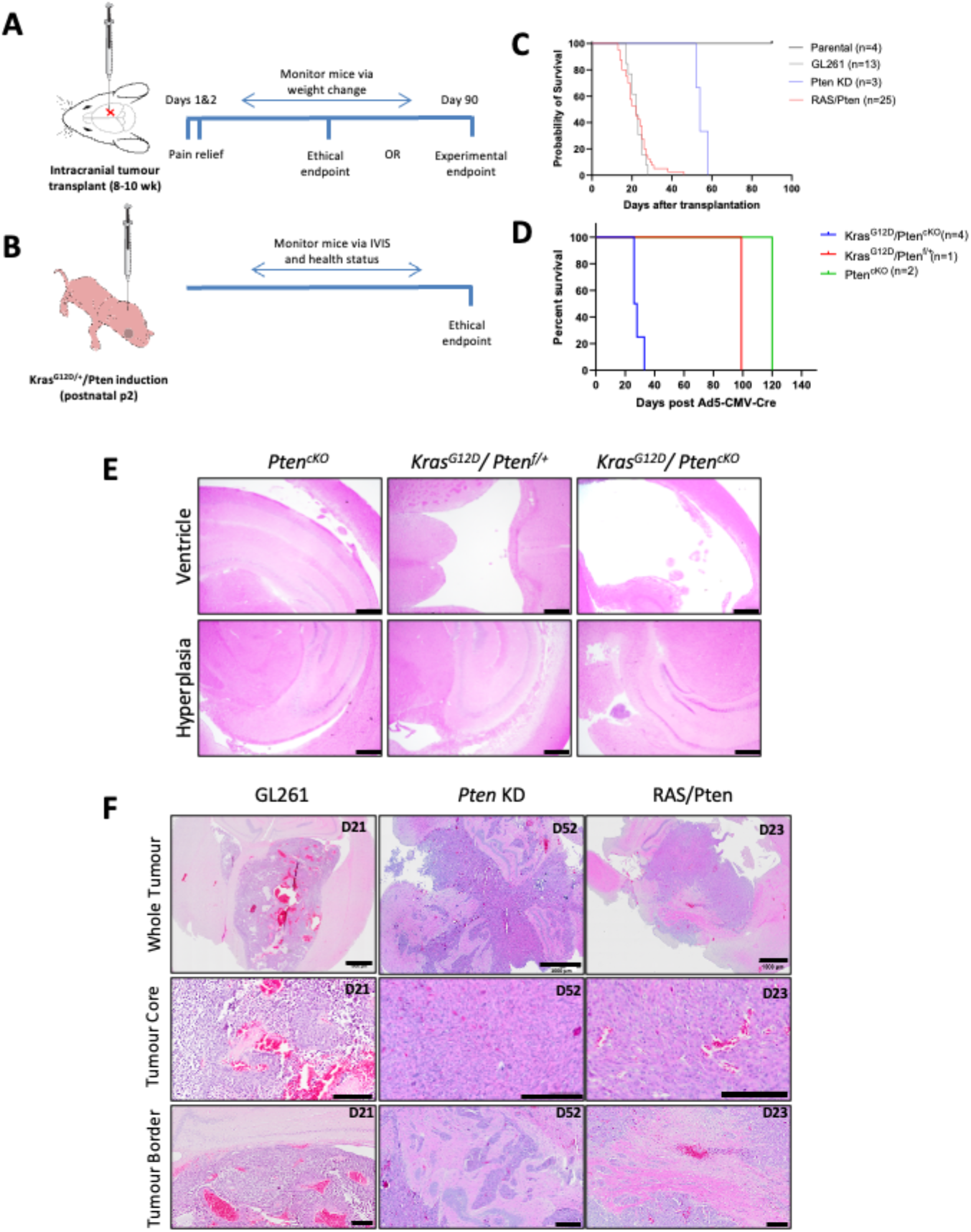
*In vivo* high-grade glioma models. A) Schematic for transplant tumor models with the parental astrocytes, Pten KD and RAS/Pten GEM-CLeM lines and GL261. B) Schematic for the generation of in vivo GEMM tumours using Ad5-CMV-Cre viral mutation system. C-D) Co-mutations to Pten and Ras induce rapid tumour formation in both transplant and GEMM systems. (C) Kaplan-Meier (KM) plot for mouse survival following cell transplantation of GL261 (grey, n=13), RAS/Pten (red, n=25), Pten KD (blue, n=3) and parental astrocytes (black, n=4). Transplantation of parental astrocytes does not induce tumour formation in up to 90 days. (D) KM plot for mouse survival following viral mutagenesis of KrasG12D/PtencKO (blue, n=4), KrasG12D/Ptenf/+ (red, n=1) and PtencKO (green, n=2). E) Virally induced Pten and Kras mutations result in small hyperplastic growth with protrusions into the ventricles. H&E-stained sections of PtencKO, KrasG12D/Ptenf/+, and KrasG12D/PtencKO tumours with representative images of ventricular areas and hyperplastic growth. F) GEM-CLeM tumours with Pten and RAS V12 mutations possess more high-grade glioma and GBM histological features than GL261 tumours. H&E-stained sections of GL261, Pten KD, and RAS/Pten tumours with representative images of the tumour cores and borders. Days post-transplant represented in each image. Unless stated, scale bars represent 200 µm.

Transplantation of parental astrocyte cells did not lead to tumor formation in 90 days, while *Pten* KD GEM-CLeM cells had symptoms indicating tumor formation at a median 54 days post-transplant (Figure 2C). Consistent with the increased proliferation seen with RAS/Pten cells *in vitro*, mice transplanted with these GEM-CLeM cells rapidly became symptomatic, with a median time to symptom onset, or survival, of 22 days. Similarly, median survival for the *Kras*^G12D^/*Pten*^cKO^/Luc GEMM was also 22-25 days (Figure 2D), with GL261 also in the same range. Differences were observed in the way animals reached ethical endpoint. In the *Pten* KD GEM-CLeM cohort, onset of neurological symptoms was the predominant endpoint (Supplemental Figure S5). Rapid onset of weight loss was characteristic of both the GL261 and RAS/Pten mice, while the GL261 mice also presented with considerable pain not seen in the RAS/Pten animals. *Kras*^G12D^/*Pten*^cKO^/Luc GEMMs displayed frequent hydrocephaly, with tumors detected using *in vivo* imaging.

Tumor histopathology at endpoint was examined. The GEMM tumors displayed hyperplastic regions protruding into the ventricles, most notable in the *Kras*^G12D^/*Pten*^cKO^/Luc mice. However, there was no evidence of substantial solid tumors in the cortex to enable comparison with the cell line models (Figure 2E). Clear differences between the GL261 and *Pten* KD derived GEM-CLeM tumors were seen on H&E staining. At day 21, GL261 tumors were characterized by large blood vessels and a spongy tumor core. Tumours showed circumscription from the surrounding brain tissue, with no signs of infiltrating tumour nuclei. Comparatively, both the *Pten* KD and RAS/Pten GEM-CLeM tumors showed significantly more cellular crowding in the tumor core than in the GL261 tumors.

Additionally, the GEM-CLeM tumors displayed some features observed in human HGG, including multinucleated giant cell morphology, and necrotic regions including foci of palisaded necrosis. The *Pten* KD and RAS/Pten tumours also showed substantial vascularisation, with a high abundance of small blood vessels present throughout the GEM-CLeM tumours. While the GL261 and GEM-CLeM tumors were more circumscribed than human HGG, both the *Pten* and RAS/Pten tumors were observed growing beyond the tumor core, aggregating as perivascular tumour nests in the surrounding vasculature (Figure 2F). These results indicated that despite similar average time to onset, the Pten KD and RAS/Pten models displayed greater histological similarities to human HGG/GBM than GL261.

### Generation of low-grade glioma GEM-CLeM lines

To generate a GEM-CLeM model of a low-grade glioma, *p53* was knocked down with CRISPR/Cas9 and human *IDH1*^R132H^ mutant protein was stably expressed (Figure 3A). CRISPR-induced indel mutation was confirmed by heteroduplex assay (Supplemental Figure 4B) and WGS (Figure 3B), and loss of p53 protein expression - both parental and *p53* KD cells were irradiated to stabilize p53, and knock-down decreased p53, before and after irradiation (Figure 3C). Cytosolic expression of IDH1^R132H^ protein was confirmed by microscopy (Figure 3D). Knockdown of *p53* significantly decreased growth rate *in vitro* compared with the parental cells. Addition of IDH1^R132H^ mutant increased the growth rate but not to the level of the parental cell line (Figure 3E). Interestingly, both the *p53* KD and IDH1^R132H^/p53 cell lines significantly increased migration in the scratch wound healing assay (Figure 3F). This indicated a potential phenotype switch to migration with the low-grade mutations.

**Figure 3.**
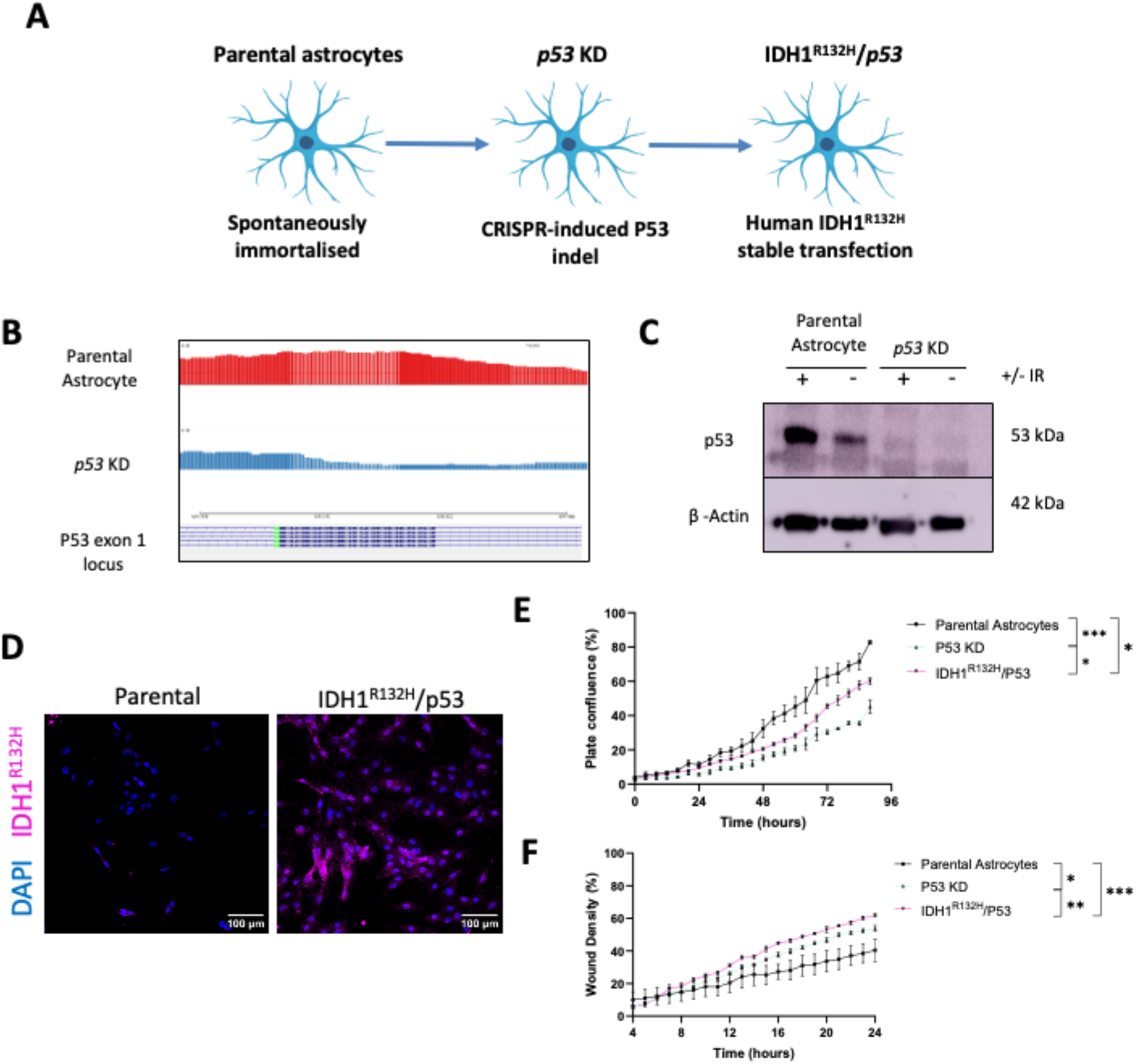
Development of the IDH1R132H/p53 GEM-CLeM cell line. A) Schematic showing the design with in vitro mutations to create p53 KD and IDH1^R132H^/p53 GEM-CLeM cell lines. B) Sashimi plot for WGS read depth over CRISPR sgRNA sites for Exon 1 of *p53* in the Parental astrocytes (red) and p53 KD (blue) sequences. C) Western blot for p53 and B-actin protein expression. Ionising radiation (+/-) with parental astrocytes and P53 KD protein lysates. B actin as a loading control. D) Immunofluorescence for human IDH1^R132H^ protein expression (pink) in the parental astrocytes and IDH1^R132H^/p53 GEM-CLeM cell line. Nuclei were stained with DAPI (blue). E) p53 KD mutation decreased proliferation rate of GEM-CLeM lines, compared with the parental astrocytes. Proliferation rate of parental astrocytes (black), p53 KD (cyan) and IDH1^R132H^/p53 (pink) lines. F) p53 and IDH1^R132H^ mutations significantly increased the rate of *in vitro* migration of the parental astrocytes. Cellular migration rate of parental astrocytes (black), p53 KD (cyan) and IDH1^R132H^/p53 (pink) lines. Student t-test, * p<0.05, *** p = 0.005-0.001. Error bars represent SEM and n = 3 biological replicates (E-F).

To investigate tumorigenesis of the low-grade astrocytoma models, 2.5×10^4^ cells of the *p53* KD and IDH1^R132H^/p53 GEM-CLeMs were intracranially transplanted, and the mice monitored for up to 90 days (Figure 4A). Consistent with the low proliferation rate observed *in vitro*, no symptoms, either neurological, pain or weight loss, were observed following transplantation of the *p53* KD derived GEM-CLeM models (Figure 4B). However, after 60 days, weight gained post-transplant had plateaued for both cohorts, but the IDH1^R132H^/p53 mice averaged lower weight than those with *p53* KD transplants (Supplemental Figure S7), suggesting tumor development. At day 90, brain tissue was collected and the transplant injection site and surrounding areas examined. Consistent with the decreased weight gain, H&E staining revealed tumor formation by the IDH1^R132H^/p53 GEM-CLeM line *in vivo* (Figure 4C). Both GFP+ tumor cell growth and cellular infiltration were observed, which were not present in the *p53* KD brain sections (Figure 4C-F). Tumor formation was supported by staining for vasculature (CD31 and CD105) and immune infiltration (CD45). At the injection site, eGFP positive GEM-CLeM cells were colocalized, with increased CD31+ blood vessels compared to surrounding brain tissue. Interestingly, this vascularisation had low CD105 expression, which indicated that while there was increased blood vessels at the tumor site, these vessels were not recently generated (Figure 4D). Throughout the eGFP+ tumor growth and in the adjacent tissue there was a high abundance of CD45+ immune cells (Figure 4E). The eGFP+ tumor mass developed surrounding an area dense with CD31+ vessels and F4/80+ macrophages. While blood vessels were prevalent in the immediately proximal tissue, these macrophages appeared to be exclusively present within the tumor mass. Conversely, TMEM119+ microglia were present in low numbers, evenly distributed in the surrounding tissue yet absent in the tumor mass (Figure 4F).

**Figure 4.**
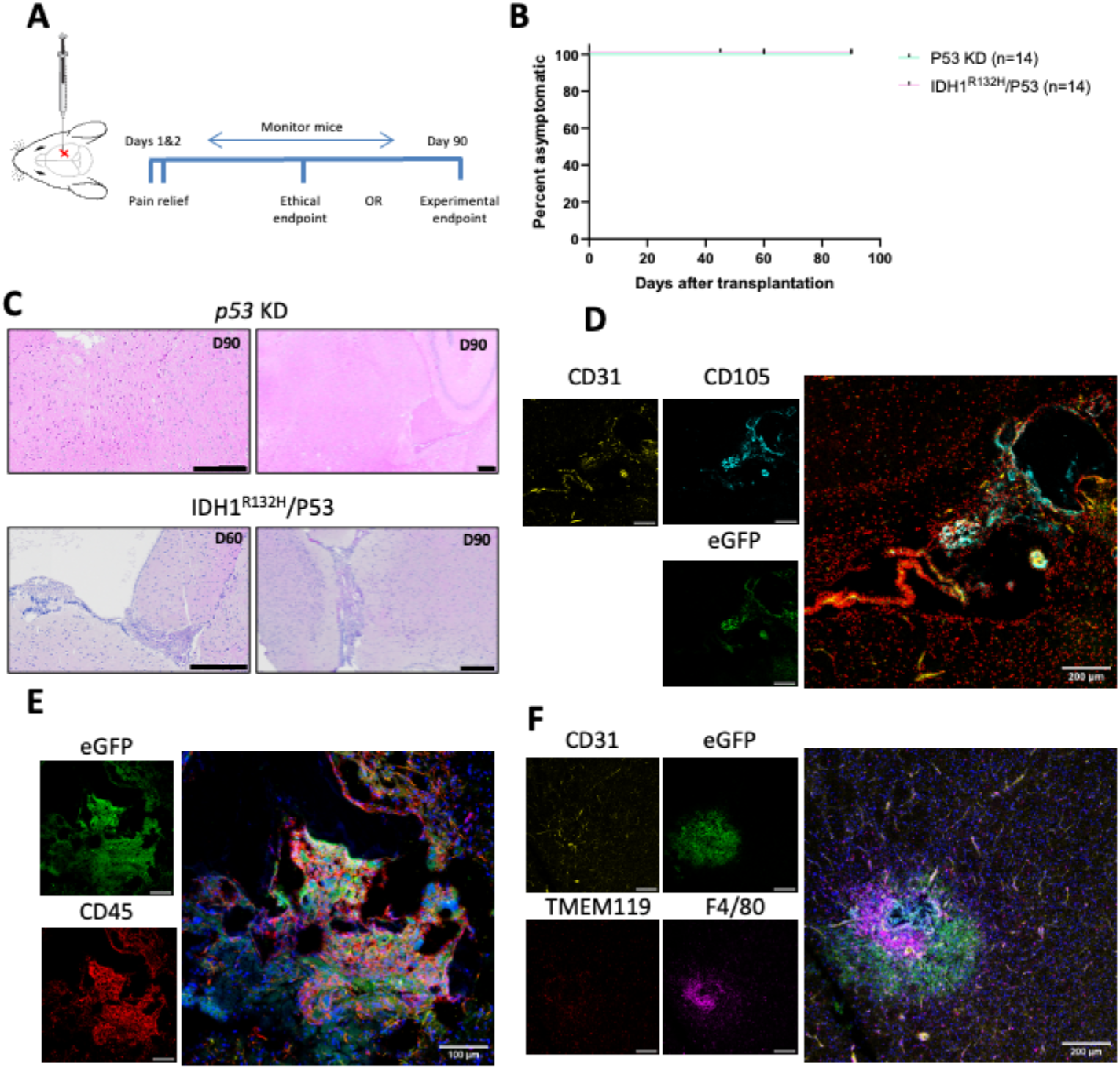
Tumor formation by the IDH1^R132H^ GEM-CLeM. A) Schematic for transplant tumour models with P53 KD and IDH1^R132H^/p53 GEM-CLeM lines. B) Mice which received transplantation of p53 KD and IDH1^R132H^/p53 GEM-CLeM lines were asymptomatic until at least 90 days. Kaplan-Meier (KM) plot for mouse survival following cell transplantation of p53 KD (cyan, n=14) and IDH1^R132H^/p53 (pink, n=14). C) Small compact tumour growth observed at transplantation site of IDH1^R132H^/p53 GEM-CLeM cells. Representative H&E-stained sections of p53 KD and IDH1^R132H^/p53 brains at the transplantation site. Days post-transplant represented in each image. D) eGFP+ IDH1^R132H^/p53 tumours are well vascularised with CD31+ blood vessels. Representative immune fluorescence (IF) image of IDH1^R132H^/p53 tumour, immunolabelled for CD31 (yellow), CD105, (cyan) and eGFP+ tumour cells (green), with nuclei stained with DRAQ7 (red). E) IDH1^R132H^/p53 tumours contained substantial immune cell infiltration. Representative IF image of immune infiltration in IDH1^R132H^/p53 tumour, immunolabelled for eGFP+ tumour cells (green) and CD45 (red). Nuclei were stained with DAPI (blue). F) Macrophage infiltration localised to IDH1^R132H^/p53 tumours tumour mass. Representative IF image of myeloid infiltration patterns in IDH1^R132H^/p53 tumour, immunolabelled for CD31 vasculature (yellow), eGFP+ tumour cells (green), TMEM119 microglia (red) and F4/80 macrophages (pink). Unless specified, error bars represent 200 µm.

### Vascularisation and immune infiltration of GEM-CLeM tumors

Microvascular proliferation (MVP) is an entity-defining histopathological feature of GBM^3^, with angiogenesis frequently an early indicator of MVP. Co-expression of CD31 and CD105 indicated the presence of newly formed endothelial vessels. In the HGG transplant models (GL261 and *Pten* KD-derived GEM-CLeM tumors), all tumors showed high levels of blood vessel formation in the tumor core (Figure 5A). Co-expression of CD31 and CD105 on these vessels indicated angiogenesis had established throughout these tumors. However, key differences in the blood vessels were apparent. GL261 tumors showed the largest vessels with wider lumen than observed in the GEM-CLeM tumours. In comparison, both *Pten* KD and RAS/Pten tumor cores had a greater number of vessels that were more compact than GL261. Consistent with the initial H&E histopathology, the *Pten* KD cells formed structures surrounding vasculature. This was seen as intense eGFP signal from areas with strong CD31+CD105+ localisation (Figure 5A). The appearance of vasculature within the *Kras*^G12D^/*Pten*^cKO^ GEMMs was different to that described in the transplant-based models. CD31+ vessels were observed in hyperplastic tissue, colocalized with highly autofluorescent red blood cells (Figure 5B). This indicated that each transplant and GEMM systems showed some degree of vascular integration.

**Figure 5.**
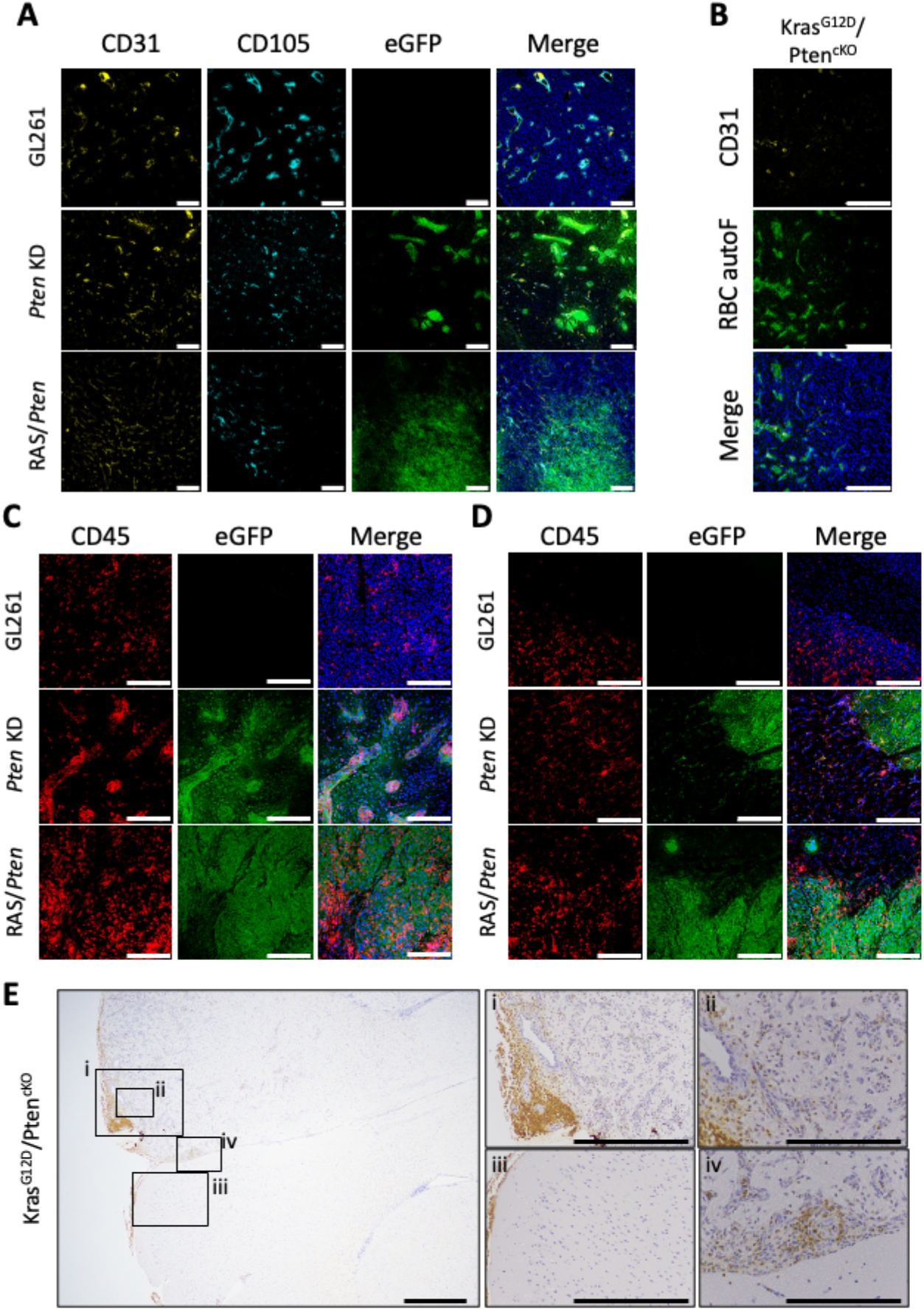
Vasculature and leukocyte distribution in high grade glioma models. A) Pten KD and RAS/Pten tumours characterised by highly abundant and small vasculature. Representative IF images of vasculature within GL261, Pten KD, and RAS/Pten high-grade glioma tumours, immunolabelled for CD31 blood vessels (yellow), CD105 blood vessels (cyan), and eGFP+ tumour cells (green). Nuclei were stained with DRAQ7 (blue). B) Large blood vessels within the hyperplastic KrasG12D/ PtencKO tumour tissue indicated by CD31+ vessels and autofluorescent red blood cell (autoF RBC). Representative IF image of blood vessels in KrasG12D/ PtencKO tumour, immunolabelled for CD31 blood vessels (yellow) and autoF RBC emission captured between 500-540 nm (green). Nuclei were stained with DAPI (blue). C-D) Higher density of immune cell infiltration within Pten KD and RAS/Pten tumours compared with GL261. Representative IF images of immune infiltration in the (C) tumour core and (D) border, of high-grade glioma models. Immunolabelled for CD45 immune cells (red) and eGFP+ tumour cells (green). Nuclei were stained with DAPI (blue). E) Asymmetric distribution of immune cells within KrasG12DD/ PtencKO brains. Representative immunohistochemistry (IHC) images stained for CD45 in KrasG12D/ PtencKO tissue with focussed regions annotated i-iv. Nuclei were counter stained blue. Unless otherwise stated, error bars represent 200 µm.

GBM tumors are characterized by a high degree of suppressive immune infiltration, leading to an immunologically cold environment dominated by macrophage and microglial infiltration^38^. To investigate the immune landscape of the GBM GEM-CLeMs, the abundance and distribution of CD45 cells was investigated in the tumor immune microenvironment (TIME). *Pten* KD and RAS/Pten tumors showed CD45 expression throughout the tumor core, and both GEM-CLeM models contained more densely packed immune infiltration than in GL261. Similarly, distribution of CD45+ cells in the *Kras*^G12D^/*Pten*^cKO^ GEMM were most abundant in areas of hyperplastic growth, although not as uniform as the transplant models. Within the adjacent brain tissue CD45+ cells were present, however, these were sparsely distributed (Figure 5E).

Consistent with previous observations^39^, GL261 immune infiltration was largely restricted to the tumor with very few CD45+ cells seen beyond the core. In the *Pten* KD tumors, CD45 expression was consistent with distribution of the vasculature. While present throughout the tumor core, the majority of CD45 infiltration into the *Pten* KD tumors was colocalized to the structures of high eGFP expression which formed around the new vessels (Figure 5C). Comparatively, a significant increase in CD45+ cells were observed outside the tumor body in the *Pten* KD and RAS/Pten sections, compared with GL261 tumors (Figure 5D). Overall, both the *Pten* KD and RAS/Pten tumours displayed the highest abundance of immune infiltration within and surrounding the tumours, most representative of the substantial immune composition of human GBM.

### RAS/Pten and GL261 tumor microenvironment comparison

Tumors that integrate malignant cells into an intact immune and stromal system are essential for glioma research. This is especially important for GBM as myeloid cells and stroma are integral to tumor growth and treatment resistance^3,9^. Fig 5 demonstrated that RAS/Pten tumors recapitulated high-grade glioma histology better than the GL261 model. To extend this, aspects of the TIME were compared between GL261 and Pten tumors.

First, the myeloid-derived innate cells; macrophage (F4/80) and microglia (TMEM119) in the RAS/Pten GEM-CLeM, GL261 and *Kras*^G12D^/*Pten*^cKO^ GEMM models were compared. Throughout all tumor sections F4/80+ cells were evenly distributed, indicating that a significant component of the infiltrating CD45 cells were monocyte derived. Interestingly, key differences in the localisation of TMEM119+ microglia were observed. GL261 tumors had localisation of TMEM119+ microglia to discrete areas, more closely aggregated than the myeloid-derived F4/80+ cells. In contrast, the RAS/Pten tumors showed substantially higher levels of F4/80+ alongside a relatively low level of TMEM119+ cells. Additionally, the microglia present in the RAS/Pten tumors were not aggregated together as observed in the GL261 sections (Figure 6A). In contrast, the macrophage and TMEM119+ cells were less abundant within the *Kras*^G12D^/*Pten*^cKO^ hyperplastic regions. F4/80+ cells were most frequently localized to the hyperplastic ventricle protrusions, while the microglia were observed throughout the adjacent tissue (Figure 6B).

**Figure 6.**
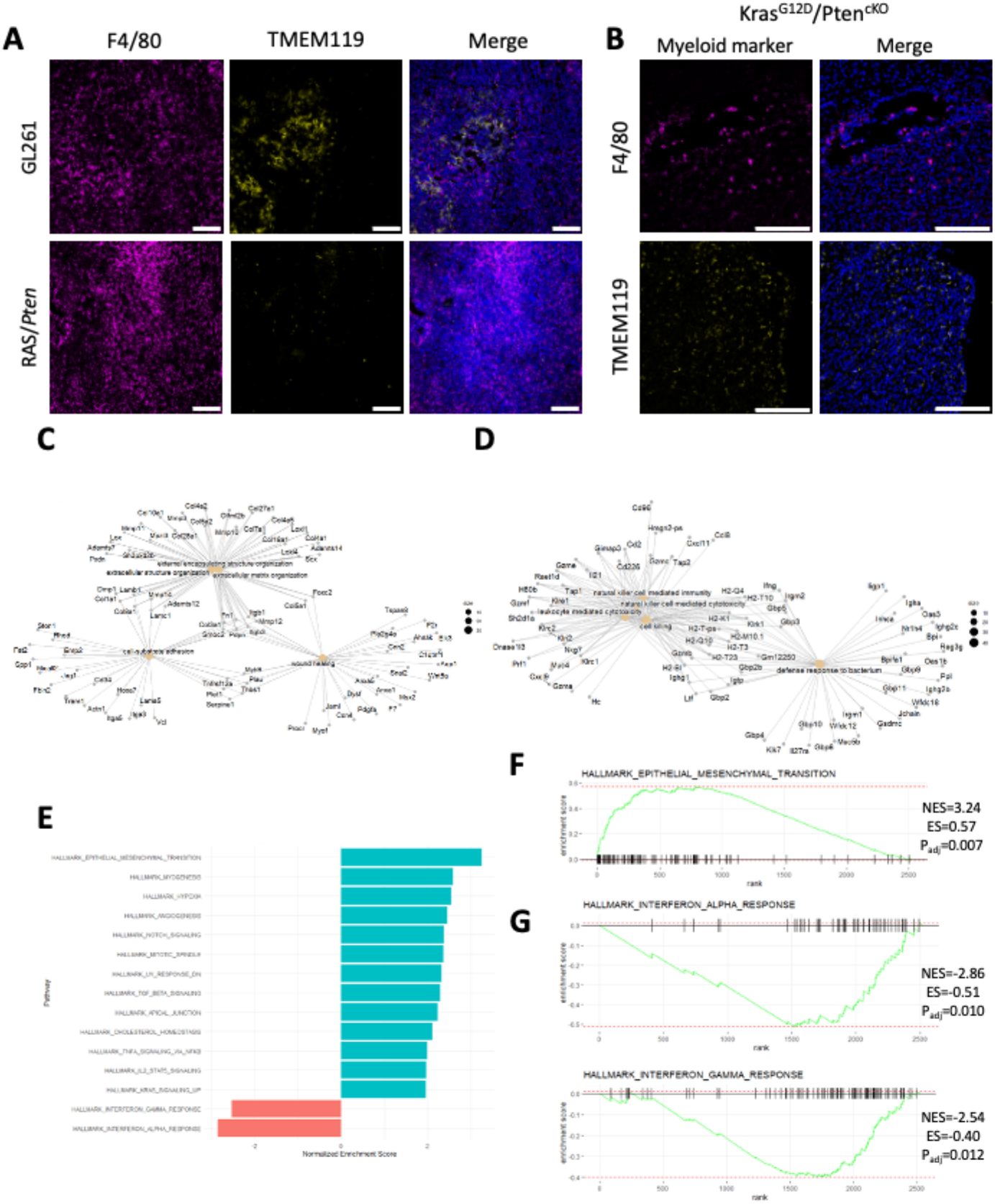
Innate immune cell infiltration and differential gene expression in glioblastoma models, A) RAS/Pten tumors dominated by macrophage infiltration, with minimal microglial, while GL261 tumours contained notable distribution of both myeloid cells. Representative immune fluorescent (IF) images of myeloid-derived cell infiltration of RAS/Pten and GL261 tumors, immunolabelled for F4/80 macrophages (pink) and TMEM119 microglia (yellow). Nuclei were stained with DAPI (blue). B) Distinct distribution trend of macrophage and microglia within KrasG12D/ PtencKO hyperplastic regions. Representative IF images of KrasG12D/ PtencKO tumors, immunolabelled for F4/80 macrophage (pink) and TMEM119 microglia. Nuclei were stained with DAPI (blue). C-D) Clear differences in GO:BP transcriptional phenotype between RAS/Pten and GL261 tumors: RAS/Pten tumours characterised by extracellular matrix (ECM) transcripts, while GL261 tumors were characterised by inflammatory immune transcripts. Gene-concept network (CNet) plots representing the expression of genes shared by upregulated gene GO:BP analysis comparing (C) RAS/Pten (n=6) and (D) GL261 (n=6) tumors. Size = number of genes in GO:BP pathway. E) GSEA shows RAS/Pten tumors enriched for multiple Cancer Hallmark gene sets corelated with GBM, while GL261 tumours enriched in inflammatory interferon gene sets. Bar plots representative of normal enrichment scores (NES) for significantly enriched (Padj < 0.05) Cancer Hallmark gene sets between Ras/Pten (cyan) and GL261 (red) tumors. F-G) Enrichment plots for the most significantly enriched Cancer Hallmark gene sets in RAS/Pten tumors.

To further characterise phenotypic differences between RAS/Pten high-grade glioma model and GL261, we performed bulk RNA-sequencing at ethical endpoint and analysed differential gene expression. Direct comparison of biological processes (GO:BP) between the two models highlighted that gene expression related to tissue and matrix remodelling was most significantly upregulated in the RAS/Pten tumor. They were strongly enriched for genes involved in ECM remodelling, including multiple isoforms of collagen (Figure 6C). In multiple cancers, including glioma, increased expression of collagen is negatively correlated with prognosis and contributes substantially to tumor aggression and invasion^40^. Additionally, the expression of other ECM proteins were also elevated in the RAS/Pten tumors, such as fibronectin (*Fn1*), podoplanin (*Pdpn*) and osteopontin (*Spp1*). These ECM-related proteins in particular reflected substantial vasculature within the tumor^41^, and a higher level of stromal cell integration compared with GL261 tumors (Supplemental Figure 8A).

In comparison, the enriched GO:BP pathways within the GL261 tumors reflected an inflamed, anti-tumor immune response. Upregulation of multiple MHCII complex genes (*H2* genes) together with interferon-gamma-related pathways indicated the presence of a more active inflammatory immune response in the GL261 tumors compared with the RAS/Pten model (Figure 6D). Further analysis indicated substantial expression of gene pathways for leukocyte and lymphocyte cell killing (Supplemental Figure 8B). Importantly, this was consistent with preclinical studies demonstrating that GL261 are highly susceptible to immune therapies, such as immune checkpoint inhibition and CAR T cells^42-44^.

Gene set enrichment analysis (GSEA) of the Cancer Hallmarks demonstrated several hallmarks associated with high-grade glioma and GBM were significantly upregulated in the RAS/Pten tumors, including epithelial-mesenchymal transition (EMT) and angiogenesis (Figure 6E). The EMT Hallmark was the most enriched gene set within the RAS/Pten tumors, reflective of the substantial number of genes associated with cellular migration and invasion (Figure 6F). This enrichment of EMT genes aligned strongly with the characterisation of mesenchymal GBM, described by Phillips et al. (2006)^5^ and Verhaak et al. (2010)^4^, and more recently the mesenchymal (MES)-like cellular phenotype described by Neftel et al. (2019)^45^. GBM tumors with a higher abundance of MES-like cells confer a worse survival due to a combination of factors such as an increased immune infiltration^4^, higher levels of radioresistance^46^ and a greater tendency to invade beyond the tumor margin^8^.

The enrichment of ‘angiogenesis’ gene set (Figure 6E) was consistent with the confocal microscopy demonstrating a greater abundance of CD105+ newly formed vasculature within the *Pten* KD derived GEM-CLeM tumors than GL261. Further GSEA utilising GO:BP pathways confirmed increased expression of vascular development and endothelial cell genes (Supplemental Figure 8). Together, these results suggested that the RAS/Pten tumors contained elevated vascular proliferation compared with GL261, in line with the presentation of human GBM.

In contrast, interferon pathways were the only significantly enriched Cancer Hallmark gene sets in the GL261 tumors (Figure 6G). Together with previous GO analyses, these results suggested a greater level of cell killing and inflammation within GL261, compared with the RAS/Pten KD tumors. This reflects the widely reported immune-responsive phenotype of the GL261 tumors^25,47^. In addition to a relative decrease in the expression of genes involved in interferon signalling, the RAS/Pten tumors showed a significant increased expression of the ‘IL2/STAT5 signalling’ and ‘TGFβ signalling’ gene sets (Figure 6E). While these pathways are complex and contribute to many different biological pathways, this upregulation of expression in these gene sets can be associated with increased immune suppression in a tumor context. In glioma, the increase in TGFβ signalling has been specifically associated with altering the TME through increased abundance of regulatory T cells^48^ and the expansion of suppressive endothelial cells^49^.

## Discussion

The GEM-CLeM represents a new approach to modelling glioma, combining the characteristic driver mutations of human disease in an immortalized murine astrocyte. The phenotype of the tumors from genetically engineered astrocytes reinforced that acquisition of mutations by a committed glial cell is an effective way to generate glioma. This does not exclude the neural stem cell as a potential cell of origin for glioma, instead it represents one way that glioma can initiate.

Only limited characterisation of the parental astrocyte line has been carried out. There were no substantive alterations to copy number observed in the WGS data of the parental line, but there will undoubtedly be changes to DNA sequence, structure or copy number associated with spontaneous immortalisation of the primary astrocyte cells. However, the parental astrocyte line will likely have a low mutation burden compared to established murine glioma cell lines such as GL261 or CT2A, which were generated by chemical mutagenesis^50^. This probable low mutation background is a particular advantage for the *IDH1*^R132H^ GEM-CLeM line. Many studies have expressed mutant *IDH1* into established GBM cell lines^51-52^, but *IDH1* mutation is an early event in LGG formation^7^. By starting from a low mutational background, the impact of IDH1 mutation can be investigated, in isolation from more aggressive driver mutations that could confound the results in other studies.

Importantly, the tumors that resulted from the IDH1^R132H^/p53 KD cells were slow-growing with distinctive immune and histological features, indicating that these mutations drive a very similar phenotype in the murine cells as in the human cells^2^.

Reassuringly, comparing the RAS/Pten GEMM and GEM-CLeM tumors demonstrated that driver mutations in the cell line led to tumors in the same timeframe as in the viral-induced GEMM model, with similar immune infiltration. However, GEM-CLeM tumors grew to a larger size in that timeframe, making them more viable for analysis of stromal and immune components of glioma. The relevance of the stromal components of the RAS/Pten tumors was highlighted by the gene expression analysis which showed that they had elevated stromal and ECM transcripts compared to GL261, particularly high expression of collagen transcript consistent with greater recapitulation human GBM histology^40^.

One use for the GEM-CLeM model is to investigate the activity of candidate genes and putative driver mutations in glioma formation and phenotype. Loss of Pten function in the GEM-CLeM was sufficient to produce proliferative, pleomorphic multi-nucleated giant cells, and tumors with palisading necrosis, an essential diagnostic criterion for GBM^6^. Addition of activated RAS significantly increased tumor growth in both the GEMM and GEM-CLeM models. However, the RAS/Pten derived tumors do not have all the characteristics of human GBM. Notably, the tumors are circumscribed, without the diffuse infiltration typical of human HGG, and there is no microvascular proliferation.

This indicates that further drivers are required for the full GBM phenotype. Both gain and loss of function mutations can readily be engineered into the RAS/Pten GEM-CLeM cells to identify the driver/s responsible for invasion, microvascular proliferation or for other characteristics of interest. Similarly, the activity of other key drivers such as EGFR or TERT can be examined, in the low mutation background, as can other genes of interest.

Another important application for the GEM-CLeM model is in immunotherapy. Immune modulating approaches for glioma and glioblastoma have not achieved the success seen in melanoma and other cancers. While there are likely multiple reasons for this, the widespread use of immunologically “hot” models like GL261, as confirmed by data presented here, undoubtedly plays a role. The presence of high numbers of immune suppressive cells in the GEM-CLeM makes them a much more realistic model to test immunotherapeutic approaches such as immune checkpoint blockade and CAR-T cells. Importantly they can be used to develop and test strategies to circumvent the intense myeloid-driven immune suppression in human glioblastoma.

In conclusion, the GEM-CLeM system is a substantial improvement on preclinical transplantation models for glioma and glioblastoma, with better representation of the stromal and immune features of human disease. The driver mutations in the transplanted cell line generate tumors very similar to a virally-induced GEMM, indicating conservation of genetic and signalling pathways. The GEM-CLeM can be used now to replace other transplantable cell lines but can also be further engineered to improve the phenotype, and customized for specific experimental questions such as characterization of new drivers, and of drug targets.

## Ethics

All mouse experiments were carried out under the approval of the animal ethics committees of Te Herenga Waka Victoria University of Wellington (VUW AEC30425 and AEC30735, and the Walter and Eliza Hall Institute of Medical Research (WEHI AEC 2022.004).

## Authorship

Experimental design – D.R.F., I.F.H., M.J.M, S.F., S.A.B., J.R.W.; Implementation – D.R.F., D.L., G.U., B.K., S.J.O., A.V.; Analysis and interpretation – D.R.F., D.L., D.E., L.S., I.F.H., S.F., J.R.W., S.A.B., M.J.M. ; Writing and review – D.R.F., D.L., S.J.O., A.V., D.E., L.S., G.U., B.K., I.F.H., S.F., J.R.W., S.A.B, M.J.M.

Data will be made available upon reasonable request.

## Acknowledgements

pX330 CRISPR plasmids targeting murine Pten and P53 were a kind gift from Tyler Jacks [Pten: Addgene_59909; P53: Addgene_59910], pcDNA3-H-Ras_V12 was a gift from Julian Downward (Addgene_39504) and pcDNA3-Flag-IDH1-R132H was a gift from Yue Xiong (Addgene_62907). The Kras^G12D^ and Pten^fl^ alleles were a kind gift from K. Sutherland, WEHI. Some of this material was presented at the annual meeting of the Society for Neuro-Oncology 2024 and at the annual meeting of the European Association of Neuro-Oncology in 2024.

