## Supplementary Figures for "A new genetically engineered transplant model of glioma recapitulates key phenotypes of low- and high-grade gliomas"

### Sup Fig 1

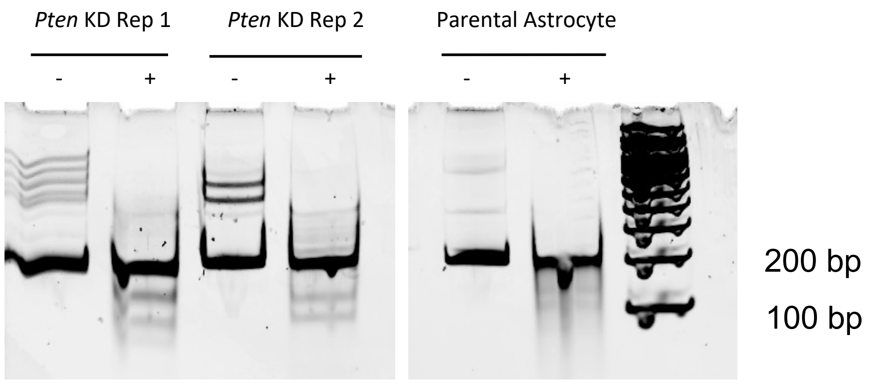


Supplementary Figure 1: Heteroduplex screening for *Pten* indel mutations. Heteroduplex assays were carried out in duplicate, Surveyor® digested (+) and non-digested (-), with parental astrocytes used as a negative control. The presence of an indel mutation in one allele was tested by formation of heteroduplex. A PCR was carried out, and product generated from both alleles was denatured and allowed to re-anneal. If each allele differed slightly due to the CRISPR-mediated generation of an indel mutation, some of the PCR product that re-annealed will be from the different alleles, and would be different lengths. This generates a small bulge in the reannealed DNA, which is recognised and cleaved by the Surveyor endonuclease. Primers were designed to flank the *Pten* sgRNA binding locus asymmetrically. The forward primer was 64 bp upstream of the CRISPR cut site while the reverse is 125 bp downstream. Therefore, if indels were present in one allele, after the digestion by Surveyor® three bands would be seen: one at 189 bp representing the PCR product which correctly reannealed; and two fragmented bands representing the 125 and 64 bp ends on either side of the sgRNA binding site.

### Sup Fig 2

***Pten* sgRNA OTS 1**

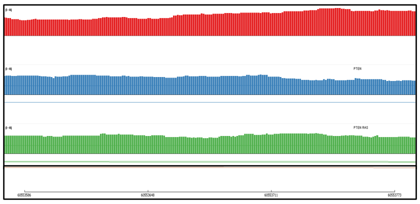

***Pten* sgRNA OTS 2**

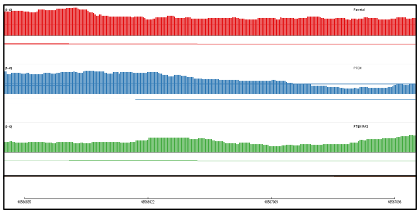

***Pten* sgRNA OTS 3**

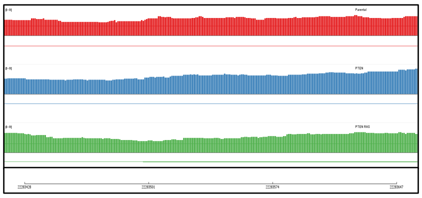

***p53* sgRNA OTS 1**

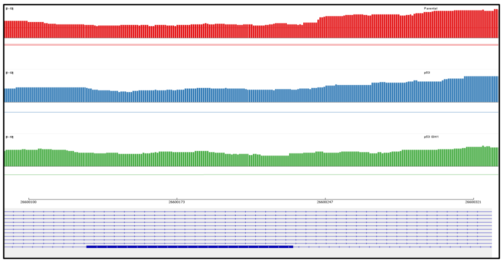

***p53* sgRNA OTS 2**

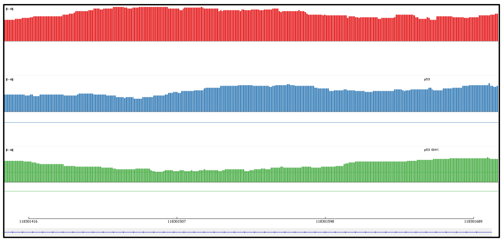

***p53* sgRNA OTS 3**

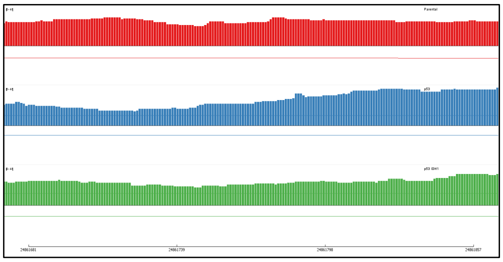

***p53* sgRNA OTS 4**

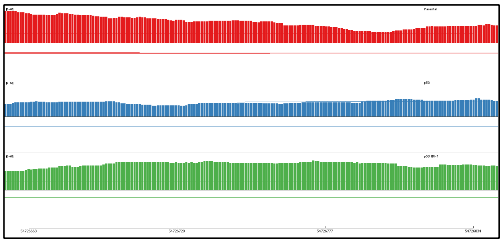

***p53* sgRNA OTS 5**

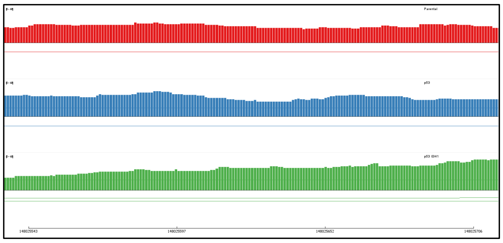

Supplementary Figure 2: Potential binding sites for the Pten (left) and *p53* (right) sgRNA were examined in the whole genome sequencing data using IGV, to determine whether there had been any off-target CRISPR activity. Red, parental DNA. Blue, *Pten* KD (left) or *p53* KD (right). Green, RAS/Pten (left) or p53/IDH1$^{R132H}$ (right).

Sup Fig 3

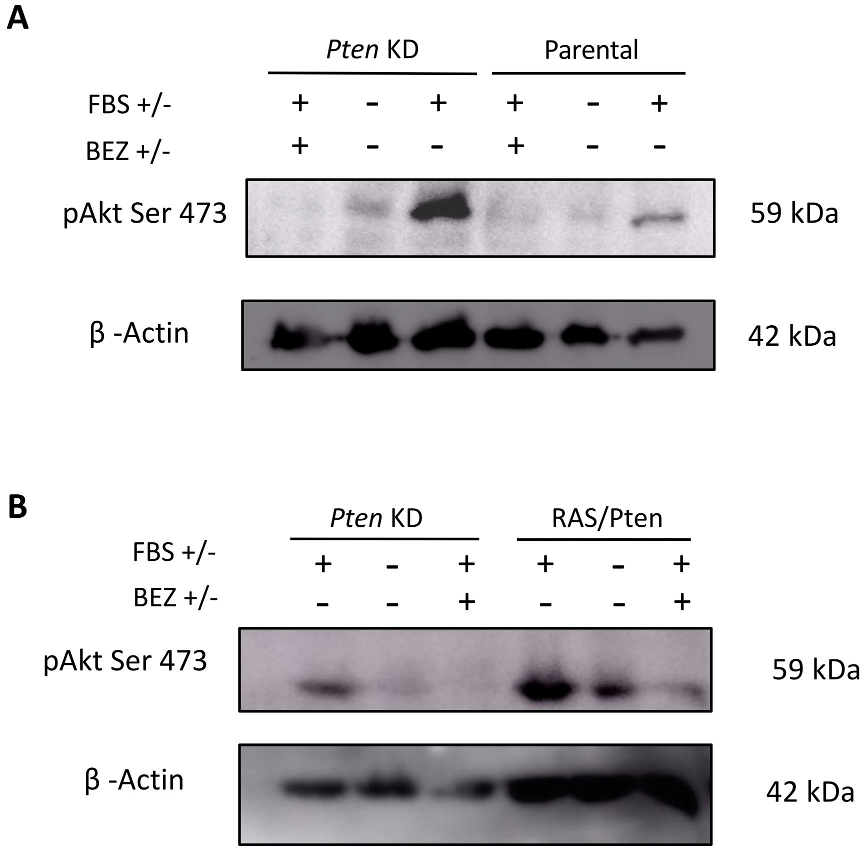


Supplementary Figure 3: Activation of Akt phosphorylation in *Pten* KD cells. Signal transduction was reduced by either overnight serum starvation (FBS-) or treatment with the PI3K/AKT inhibitor BEZ-325. Serum starved cells were then stimulated with FBS, and cells collected and lysed after 15 minutes. Cells were treated as described, then the lysate blotted for A. *Pten* KD cells and parental cells. B. *Pten* KD and RAS/Pten cells.

### Sup Fig 4

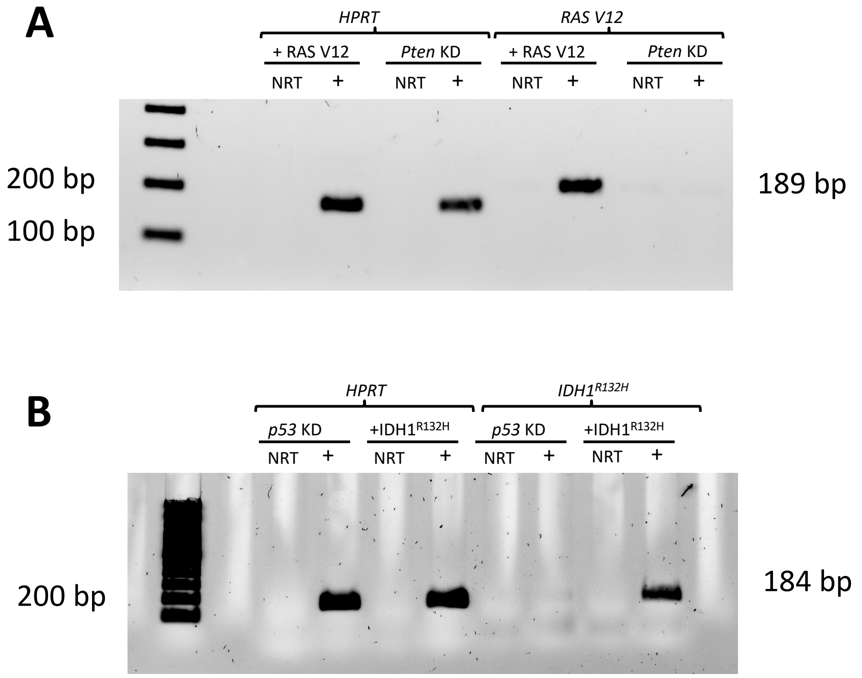


Supplementary Figure 4: RT-PCR confirmation of oncogene expression after stable transfection. RT-PCR was used to detect transcript specific for (A) RAS V12 (top, 189 base pairs) and (B) $IDH1^{R132H}$ (bottom, 184 base pairs) expression. HPRT was used as a positive control (left side of each gel), and a "no reverse transcriptase" control (NRT) was carried out for each sample. Both RASV12 and $IDH1^{R132H}$ were only detected in the transfected cells. Primer sequences were: RAS V12, 5' GGAAGCAGGTGGTCATTGAT-3 and 5'-ACGTCATCCGAGTCCTTCA-3'; $IDH1^{R132H}$, 5'-AAAAATCAGTGGCGGTTCTG-3' and 5'-TATAGCTTCTGCGGCATCCT-3'; HPRT, Quantitect (QIAGEN).

Sup Fig 5

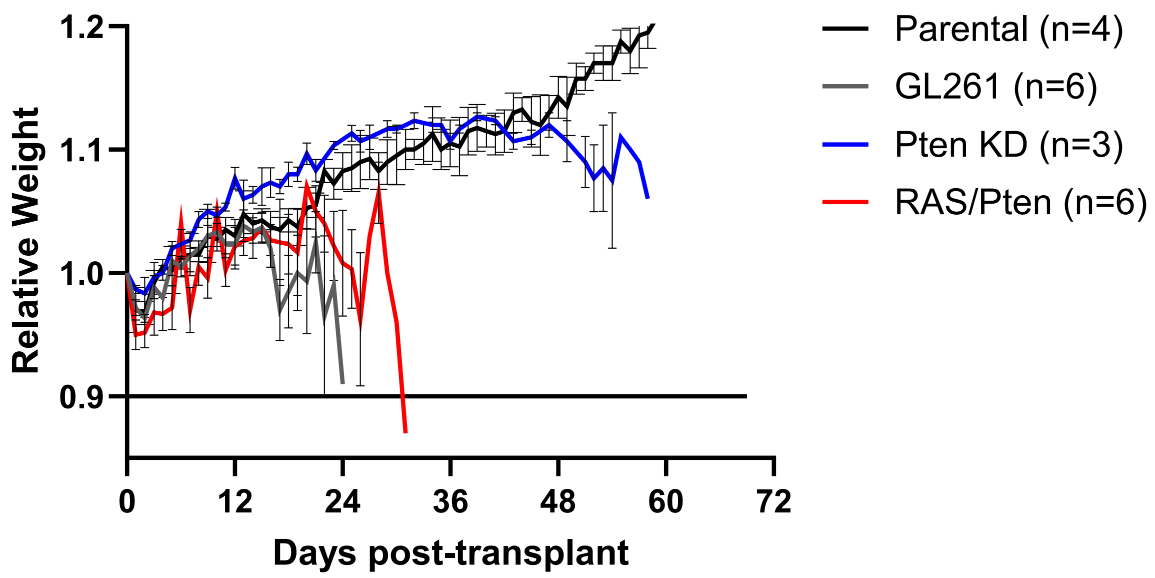


Supplementary Figure 5: Average mouse weights, relative to the date of cell transplant (day 0). The line at 0.9 indicates the ethical endpoint of 10% weight loss. Mice with GL261 (grey, n=6) and RAS/Pten (red, n=6) transplants had rapid weight loss, while mice with Pten KD cells (blue, n=3) slowly lost weight. Mice with parental cells transplanted (black, n=4) continued to gain weight. Error bars represent SEM.

### Sup Fig 6

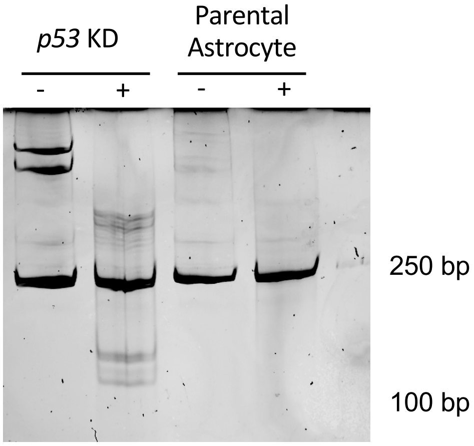


Supplementary Figure 6: Heteroduplex screening for p53 indel mutations. Heteroduplex assays were carried out in duplicate, Surveyor® digested (+) and non-digested (-), with parental astrocytes used as a negative control. The presence of an indel mutation in one allele was tested by formation of heteroduplex. A PCR was carried out, and product generated from both alleles was denatured and allowed to re-anneal. If each allele differed slightly due to the CRISPR-mediated generation of an indel mutation, some of the PCR product that re-annealed will be from the different alleles, and would be different lengths. This generates a small bulge in the reannealed DNA, which is recognised and cleaved by the Surveyor endonuclease. Primers were designed to flank the p53 sgRNA binding locus asymmetrically. resulting in a predicted result of the PCR product at 288 bp, and two fragmented bands at 135 bp and 153 bp respectively.

Sup Fig 7

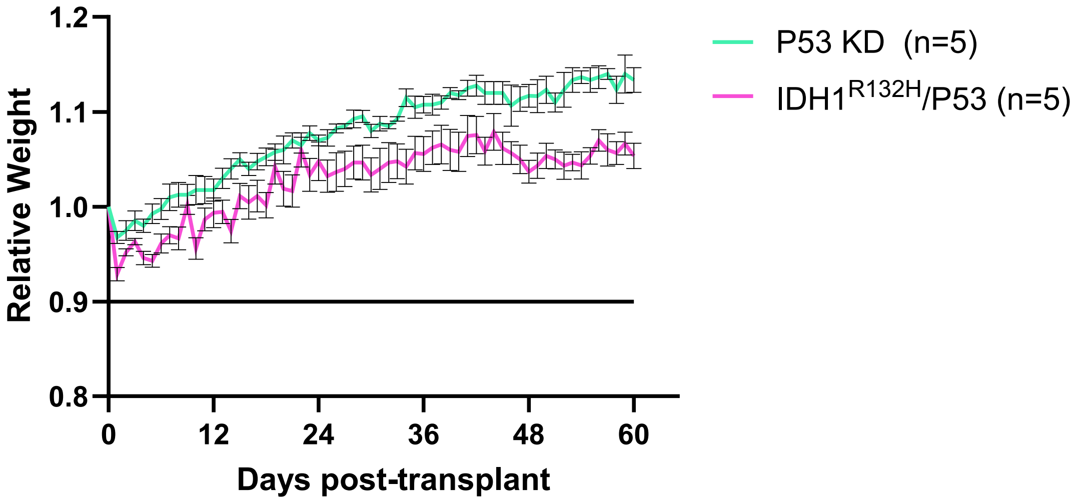


Supplementary Figure 7: Average mouse weights, relative to the date of cell transplant (day 0). The line at 0.9 indicates the ethical endpoint of 10% weight loss. Mice with *p53* KD (green) and IDH1$^{R132H}$/p53 (pink) transplants did not lose weight over 60 days, but the mice with IDH1$^{R132H}$/p53 transplants plateaued at a lower weight. Error bars represent SEM

### Sup Fig 8

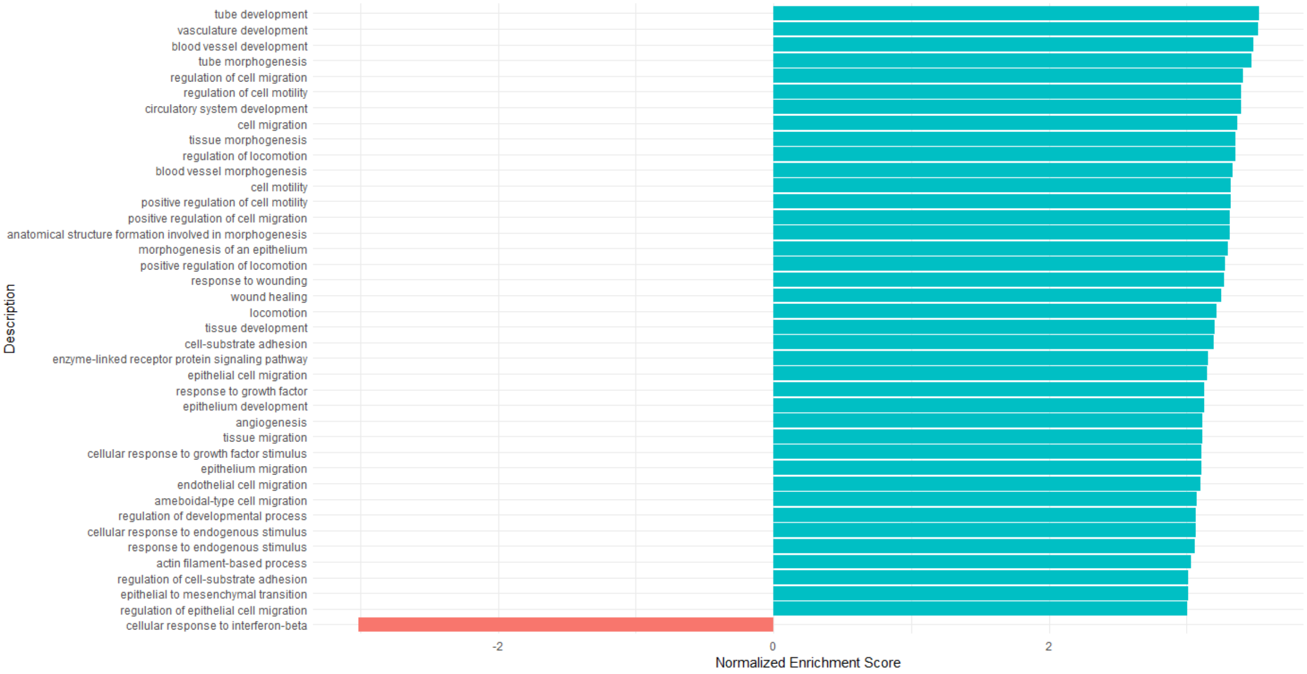


Supplementary Figure 8: RAS/Pten tumors display an increase in vascular development and a decrease in interferon production compared with GL261**.** Gene set enrichment analysis (GSEA) for gene ontology: biological processes (GO:BP) gene sets, comparing differentially expressed genes (DEGs) between RAS/Pten (n=6) and GL261 tumors (n=6). Bars represent significant GO:BP gene sets (adjusted p value < 0.05). GSEA results shown as normalised enrichment score (NES); positive NES (cyan; gene sets with NES > 3) = upregulated in GL261, negative NES (red; gene sets with NES < -3) = upregulated in GL261 tumors.
